# A generalized quantitative antibody homeostasis model: antigen saturation, natural antibodies and a quantitative antibody network

**DOI:** 10.1101/065748

**Authors:** J́zsef Prechl

**Affiliations:** Diagnosticum zrt, 1046 Attila út 126. Budapest, Hungary; MTA-ELTE Immunology Research Group, at Eötvös Loránd University, 1113 Pázmány Péter s. 1/C, Budapest, Hungary

**Author notes:** Grant information The author(s) declared that no grants were involved in supporting this work.

**Keywords:** antibody, antigen, thymus independent, thymus dependent, natural antibody, network, affinity, concentration

## Abstract

In a pair of articles we present a generalized quantitative model for the homeostatic function of clonal humoral immune system. In this second paper we describe how antibody production controls the saturation of antigens and the network of antibody interactions that emerges in the epitome space with the establishment of the immune system.

Efficient control of antigens, be it self or foreign, requires the maintenance of antibody concentrations that saturate antigen to relevant levels. Simple calculations suggest that the observed diverse recognition of antigens by natural antibodies is only possible by cross-reactivity whereby particular clones of antibodies bind to diverse targets and shared recognition of particular antigens by multiple antibody clones contribute to the maintenance of antigen control. We also argue that natural antibodies are none else than the result of thymus independent responses against immunological self. We interpret and explain antibody production and function in a virtual molecular interaction space and as a network of interactions. Indeed the general quantitative (GQM) model we propose is in agreement with earlier models, confirms some assumptions and presumably provides the theoretical basis for the construction of a real antibody network using sequence and database data.

## The GQM applied to antibody homeostasis

By definition antigens are molecules recognized by antibodies. Most definitions however fail to further elaborate what exactly is meant by recognition. The strength of the interaction between the antigen binding site (paratope) of an antibody and the antibody binding site (epitope) of the antigen is characterized by affinity, kinetics of association and dissociation and binding energy. Antibodies often recognize more than one target. Immunological assays usually require the titration of the antibody, which is the identification of lowest concentration that binds to the nominal target but does not bind to others. This is quite logical for antibodies intentionally produced in animals, but how we define the target of an antibody in vivo? By changing the concentration of antigen and antibody, saturation of any can be achieved even when affinity of the interaction is low. The absolute and relative concentration of antigens and antibodies does matter and our GQM attempts to reveal antibody function by addressing these factors.

The general equation defining equilibrium dissociation constant K_D_:

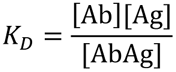

tells us that K_D_=[Ab] when [Ag]=[AbAg]. That is when antigen is half saturated, free antibody concentration is equal to K_D_. For the sake of simplicity we will regard [Ab] as the concentration of paratope and [Ag] as the concentration of epitope and we shall use the term apparent affinity to indicate that factors like multiple binding sites modulate the observed strength of the interaction. Assuming that antibodies are produced with the intent of regulating antigen availability, best control over antigen concentration is achieved when the concentration of antibody is close to the K_D_ (Fig.1). In our map this zone for a range of [Ab] and K_D_ values is defined by a line where [Ab] =K_D_, which is the line representing 50% saturation of the antigen (Fig 1). By lowering or increasing antibody production the host can release or capture antigens, and likewise by changing the efficiency of Ab binding the host can modulate antigen saturation (Fig.1). Various immunological mechanisms are responsible for removing antibody-antigen complexes, called immune complexes, from the circulation.

**Figure 1.**
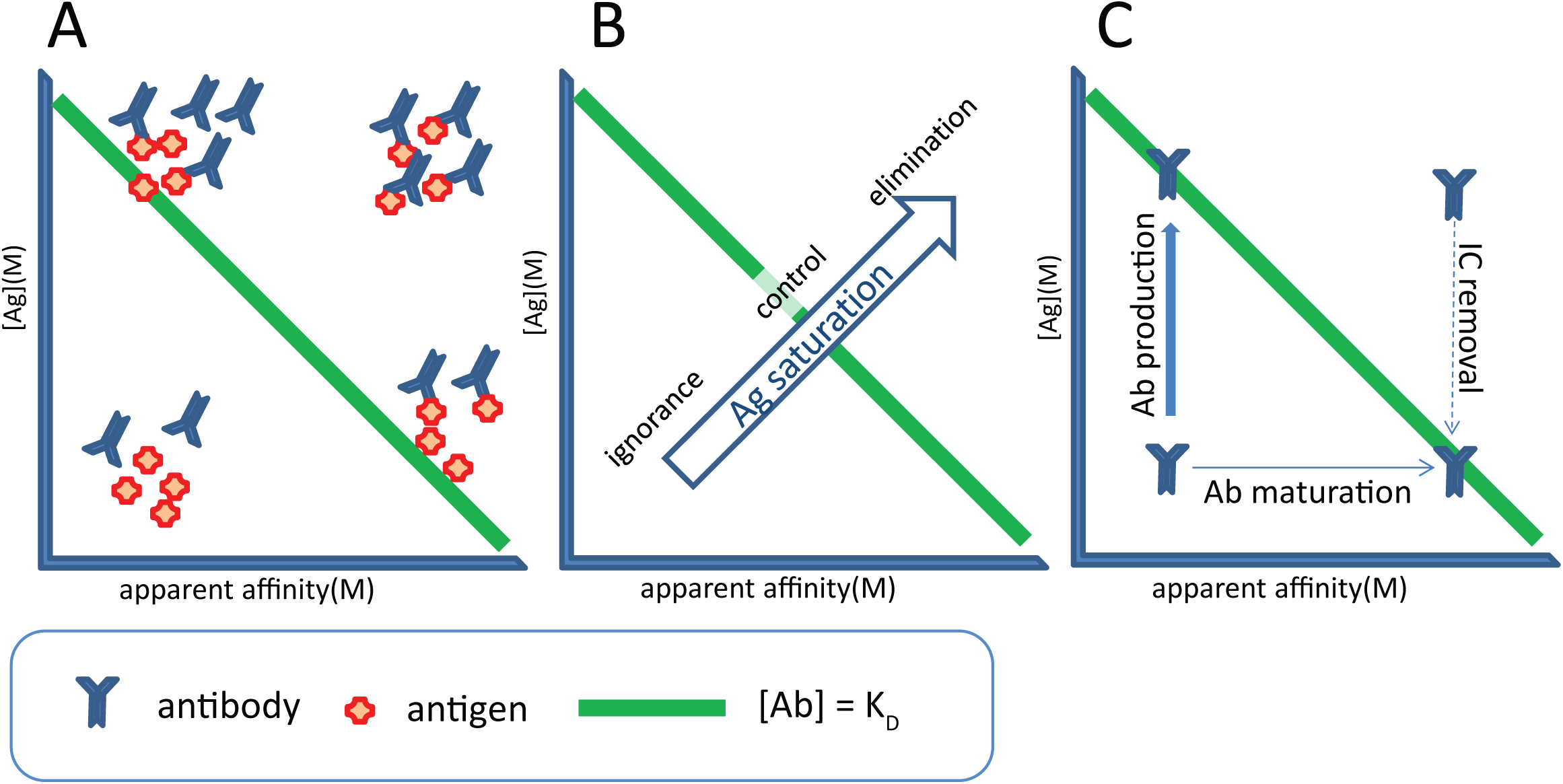
Outlines of the GQM for regulation of antibody production. A, Antibodies will saturate antigen by increasing their concentration or by increasing apparent affinity. B, Low concentrations of low-affinity antibodies do not bind antigen at relevant extent, antigen concentrations can be best controlled at around 50% saturation, while elimination is achieved by increasing saturation further. C, ASC produce antibodies to increase concentration, while prior differentiation account for increased affinity or ability to remove antigen. Immune complexes are removed by different cells and silent or proinflammatory events.

The range of [Ab] values we will be using in our model reflect actual immunoglobulin concentrations in blood plasma, and start around the concentration that a single plasma cell could achieve by continuous secretion of antibody. The range of K_D_ values includes affinity constants usually observed^1^ for antibody-antigen interactions (of K_D_=10^−6^–10^−10^ M) but extends to both lower and higher values to provide flexibility for interpreting apparent affinities. Please note that these are exactly the same dimensions, which we use in our accompanying sister paper on B-cell development (Prechl, submitted). Let us now analyze various characteristic immune responses in the order of increasing antibody-antigen interaction affinity. We will consider a single fluidic compartment, the blood plasma for this theoretical framework, however, with proper adjustments the model can be possibly extended to include the extracellular space and mucosal surfaces – sites of key importance for immunological action.

## Natural antibodies and TI antibody responses

Can low affinity antibodies mediate any relevant biological effect at all? For an antibody with K_D_=10^−6^ M a concentration of 10^−6^ M should be reached for substantial binding to its target, which is quite close to the total immunoglobulin concentration in plasma (Fig.2). Obviously, no single antibody can dominate to such an extent (except for pathological antibodies in disease, like monoclonal gammopathies). Multiple antibody binding sites on the antigen increase the apparent affinity of the interaction by avidity effects, but not to the value required here. Most likely a combination of these effects, large cumulative concentration achieved by a large number of cross-reacting antibody producing B-cell clones and avidity might confer effector functions to low-affinity antibodies.

**Figure 2.**
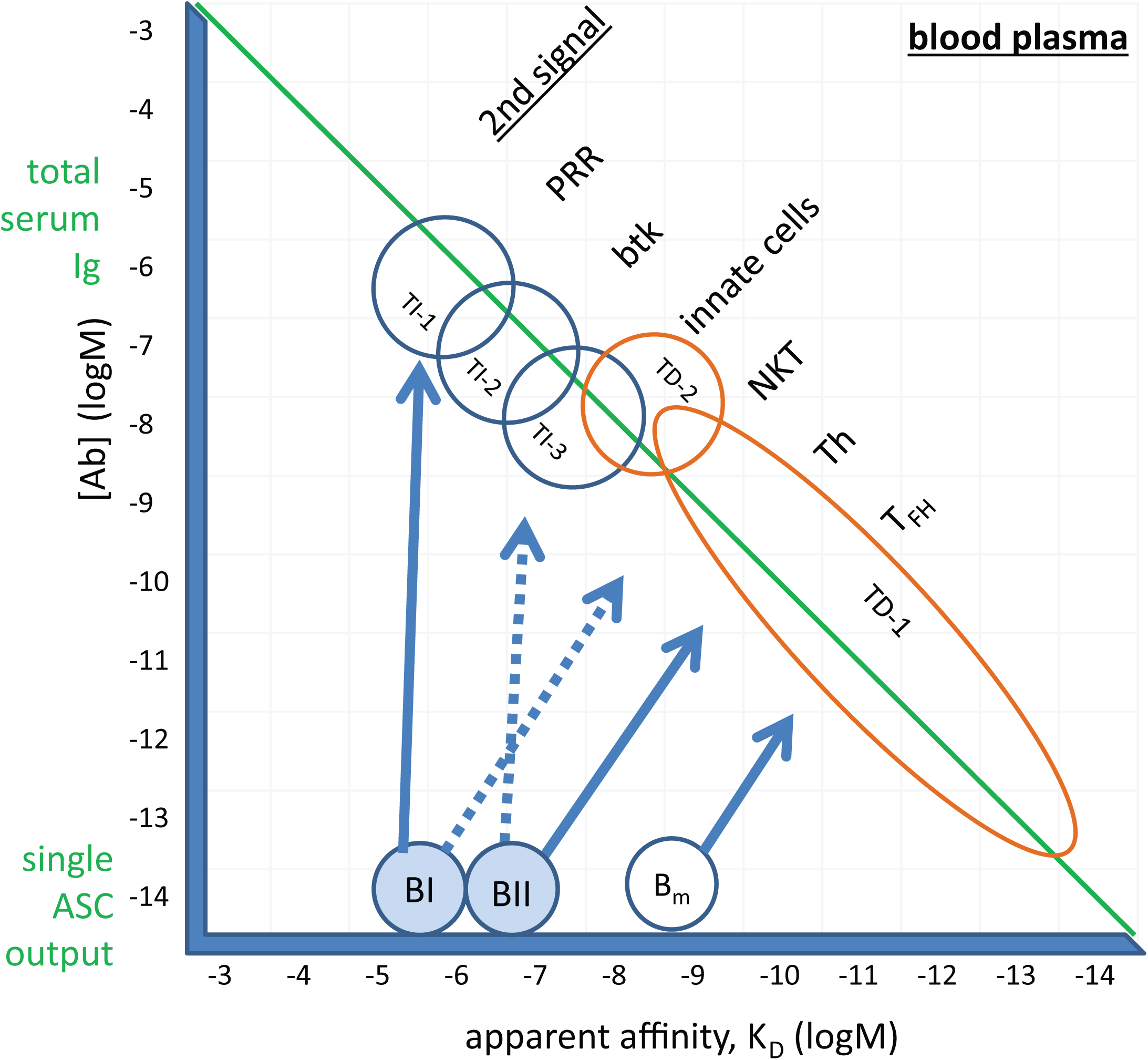
Balance of Ab and Ag achieved by different humoral immune responses. The epitope-antibody interaction landscape as defined by free antibody concentration and affinity. Second signals required by B cells for becoming antibody secreting cells are listed next to the type of immune response. The range of total serum immunoglobulin concentration and the concentration achieved by a single ASC clone are indicated. PRR, pattern recognition receptors; btk, Bruton’s tyrosine kinase; NKT, NK T cells, Th, helper T cell, T_FH_, follicular helper T cells; B_m_, memory B cell; TI, thymus independent; TD, thymus dependent

Natural antibodies are low-affinity antibodies, constitutively produced by B1 cell populations that are relatively well characterized in the mouse^2^ but only recently described in humans^3^. Natural antibodies are polyreactive or polyspecific, binding to structurally different self and microbial targets as well^4,5^, these targeted epitopes being mostly non-protein molecules. The affinity of natural antibodies to monovalent glycan has been determined to be in the range of 10^−4^ to 10^−6^ M^6^. While mostly of the IgM class^7^, functionally similar antibodies belonging to the IgA and IgG classes are also found. Besides providing immediate protection against invading microbial agents, natural antibodies are known to play key roles in the clearance of self molecules, thereby contributing to homeostatic control, suppressing inflammation and autoimmunity^8,9^. In humans B1 cells represent from less than 1 to 9 % of the circulating B cells^10^. Calculating with an average 4×10^9^ white blood cells per liter of blood, 5% B cells in white blood cells, and an average 5% of B1 of all B cells we arrive at an averaged 5×10^7^ B1 cells in 5 liters of blood, capable of a dominant contribution to the plasma antibody pool. The number of circulating B1 cells indeed shows correlation with serum IgM levels^11^.

As we have discussed in our accompanying article, our GQM assumes that B1 cells develop from immature B cells in blood, as a result of continuous BCR signaling triggered by blood borne antigens. Their precursors, immature B cells were selected based on their polyreactive property, tested by self-antigen displayed on the developing cells. Polyreactivity against common, shared molecular targets allows clusters of antibodies to cooperatively bind to highly abundant self-antigens, reaching relevant fractional saturation despite the relatively low affinity of the interactions. Any particular antibody can belong to several different such clusters, thereby increasing the concentration of antibodies against that particular antigen. In other words, **by shared, distributed recognition several different antigenic targets can be saturated by the same amount of antibody**. Cross-reactivity increases the apparent clonal diversity, because clusters of the same network respond to different targets. Removal of cellular debris, which contains a huge number of different molecules may become possible this way by a limited number of specificities and antibodies. Polyspecific, low-affinity, high off-rate interactions in the plasma result in dynamic short-lived contacts, these natural antibodies acting like a lubricant rather than like cement. On the other hand molecular aging, often presenting as aggregation, results in polymerization of the target, increasing the avidity of interactions with natural antibodies, and aiding removal. This removal process is continuous and silent. Natural IgM fixes C1q, which in turn is captured and taken up by phagocytes throughout the body, utilizing different C1q receptors^12–15^. Natural antibodies may also cover and seal leaks in the endothelium, promoting regeneration and healing^16^.

In addition to B1 cells, MZ B cells also contribute to the fast production of upon challenge^17^. These two populations are the cells responsible for thymus independent (TI) responses. Is there, then, a difference between natural antibodies and TI antibody responses? We believe there is not. Natural antibodies are defined as being produced in the absence of a known antigenic stimulus. This is possibly a wrong interpretation of the events. Accepting that B cell development involves selection of low-affinity, polyspecific self-reactivity, the antigenic stimulus is self. Natural IgM antibodies are also present in germ-free mice^18^, where the only template for antibody production is self.

**Natural antibodies are born silently and are eliminated silently**, because no inflammatory events accompany either induction of antibody production or removal of forming immune complexes. They **function to aid the catabolism of cellular debris, of aging proteins, of endothelial cells and contribute to the molecular and cellular regeneration of blood and blood vessels**. Since the same type of cells produce natural antibodies and TI antibodies, we consider these antibodies essentially the same, especially for IgM. Whereas the continuous presence of stimulatory concentrations of blood borne antigen drives antibody production by B1 cells, TI responses are elicited by further increasing Ab production and recruitment of natural memory cells, the MZ B cells. This can be triggered by administration of a proper antigen: pattern recognition receptor ligands for TI-1^19^, highly repetitive microbial and viral motifs for TI-2^20^, and recruitment of myeloid cell help for TI-3^21^ (Fig.2). Second signals provided by microbial products and cytokines can induce class switching, leading to the generation of IgA and IgG (mainly IgG1 and IgG2) antibodies. Overall, TI responses are the manifestation of antibody secreting effector function of B cells selected in the bone marrow and primed in the periphery for controlling abundant self and non-self antigens. In short, **antibodies produced by TI responses constitute the natural antibody repertoire** (Fig 3).

**Figure 3.**
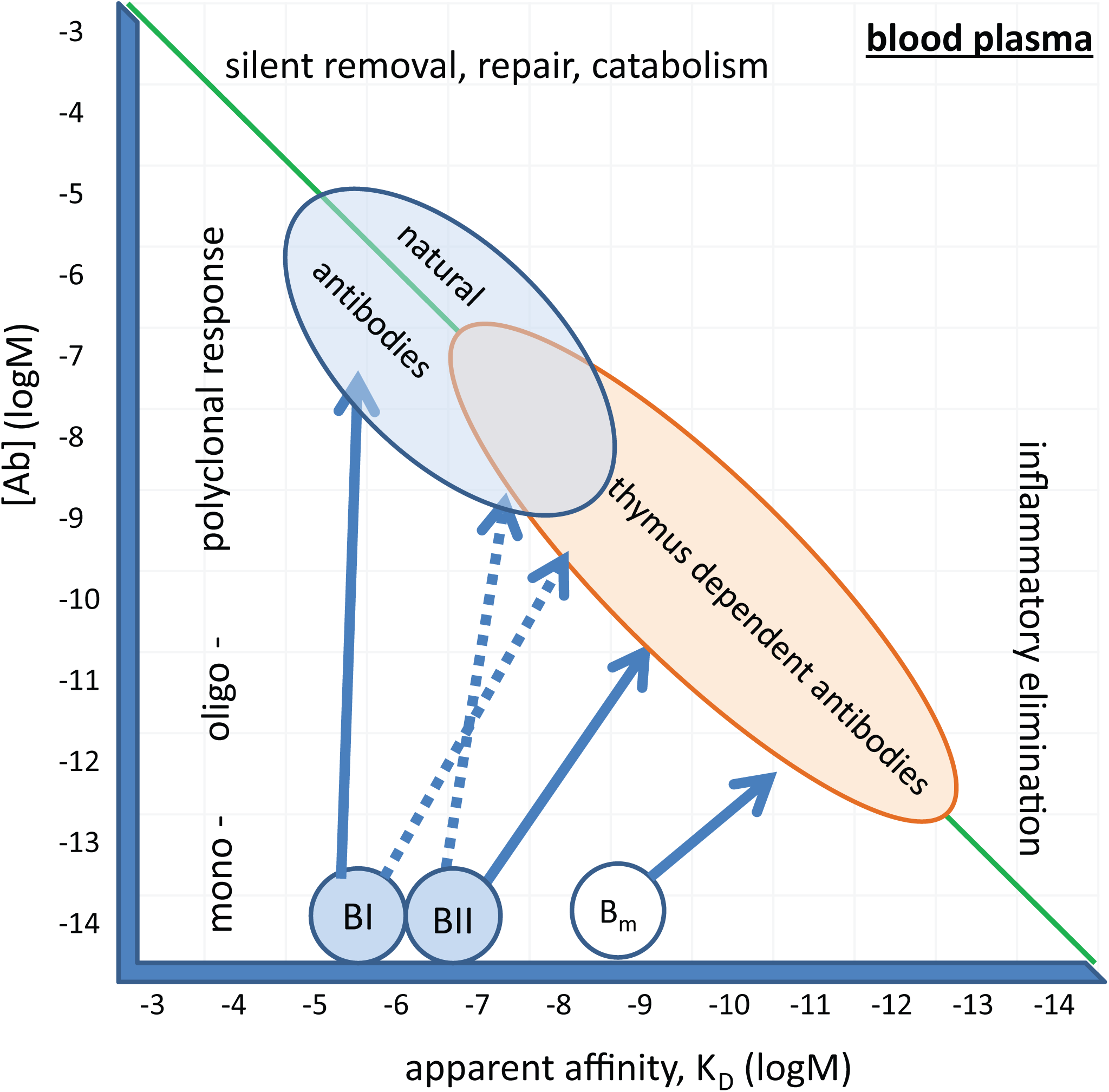
Thymus independent and thymus dependent antibody responses in the interaction landscape. Relative concentration of epitopes, antibodies and affinity of the interactions define natural and thymus dependent antibodies.

## TD antibody responses

The affinity of a given antibody to a given antigen can only be increased by changing its structure and sequence. While structural changes, such as oligomerization (IgM pentamer > hexamer, IgA monomer > dimer conversions) play role in TI responses, template driven maturation of affinity by sequence modification requires T-cell help. B cells with increased affinity to the eliciting antigen emerge in germinal center reactions, giving rise to antibody secreting effector cells (ASC) and memory B cells. Our GQM postulates that antibody secretion is an effort to decrease [Ag] and return B cells to their comfort BCR signaling zone (Prechl, submitted). These ASC leave the germinal center and circulate in blood, and are found in blood rich compartments of the bone marrow and spleen, some settling and forming long-lived plasma cells in suitable microenvironments^22^. Antibody secretion by ASC contributes to the pool of antibodies in blood. Along with the increase in affinity against the target antigen antibodies lose their polyreactivity, forming less flexible but more precisely fitting hypervariable loops to contact antigen. Repeated exposure to the same antigen (booster injections or hypervaccination) recruits memory B cells which have an already high affinity, making further steps along the affinity axis in our map (Fig.2) until limits are reached^23^. Increasing the affinity means that less and less [Ab] is required for saturating the antigen, which translates into less and less ASC required for eliciting elimination of antigen. This is the idea behind most of our prophylactic vaccines^24^. Our map suggests that even with high affinity binding of 10^−10^ M a huge number of ASC clones are needed for conferring such protection. The situation drastically changes of course, if we look at local concentrations instead of plasma concentration, where even a single ASC can generate adequate amounts of antibody.

Affinity maturation is usually accompanied by class switch, so the heavy chain class of high affinity antibodies changes from IgM to IgA, IgG and IgE. This has very important bearing on the way how immune complexes forming from these antibodies are eliminated. Complement activation and FcR-mediated effector function properties of these antibody classes can be significantly different from those of natural antibodies^25–27^. Increased affinity, mostly accompanied by decreased dissociation rates^23^, lends rigidity to the forming immune complexes, also making them more prone to elimination. Saturation by high affinity antibody means close to continuous presence of bound antibody, which is poised to lead to interaction of the complex with other components of the immune system, such as cells bearing Ig receptors or complement proteins. All **characteristics of a TD response imply that the immune system does not accept the targeted antigen** as part of the immunological self and it concerts its efforts to reject the antigen.

The properties of the BCR and antibodies are summarized in Figure 4, layers representing stages of development and antigen recognition. Immunological self is a part of the antigenome recognized as part of the host. Extended immunological self incorporates embryonic or genetic self and the molecular environment recognized by natural antibodies. Catabolism of the repertoire of molecules in the extended self or immunological self is provided for by natural antibodies. Accepted environment is the border of immunological self, with enhanced removal of the forming immune complexes. Rejected environment is recognized by high affinity antibodies which elicit strong effector functions and attempt to eliminate the antigen.

**Figure 4.**
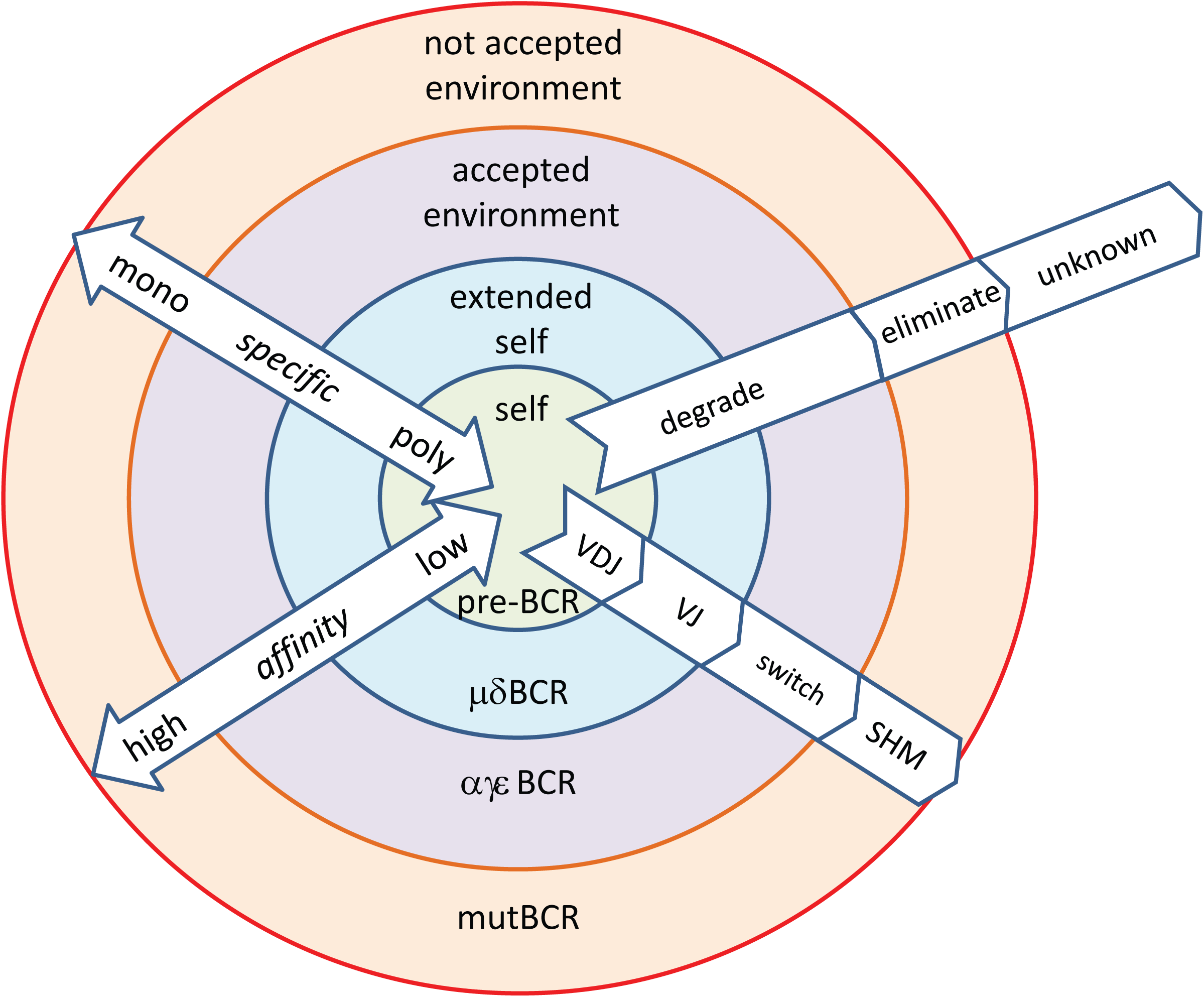
Layers of interactions of antibodies with immunological self and non-self. mutBCR, BCR with somatic hypermutation

Using the dimensions [Ag], [Ab] and K_D_ we have been able to draw a raw general map of clonal humoral immune responses, positioning B-cells at different stages of development and antibodies produced by them on these maps. With these quantitative descriptors we can now try to design a network that models immune function.

## Interpreting the humoral immune response as a network phenomenon

Let us imagine a space of epitopes. In this space certain coordinates identify interactions with the immune system. The space is fluid, epitopes can change their positions. The developing immune system seeds this space with interactions by generating antibodies that bind to epitopes. Each formed B cell with an antigen receptor (BCR) is represented as a point in this space. This will be the epitope that is recognized with the highest affinity by that particular BCR. Let us call this the cognate epitope of the antibody. Because developing fetal B cells only come into contact with self, this will be a self epitope in the epitope space of all epitopes, the epitome. Please note that the exact identity of cognate antigen can change, as the immune system encounters new epitopes, with potentially higher binding affinity. The antibodies these cells (or their effector forms) secrete will bind to this cognate epitope but also to closely related epitopes, even though with lower affinity. To characterize the collection of epitopes in the neighboring epitope space to which these antibodies bind we can now use a sphere. The amount of antibody available for binding to epitopes can be symbolized as [Ab]/K_D_, an expression called antigen binding capacity. So we can draw a sphere with a radius [Ab]/K_D_ (Fig.5). These points, representing B cells with identical antigen binding properties, and spheres, representing the capacity of antibodies to bind to the contained epitopes, will be the nodes in our network.

**Figure 5.**
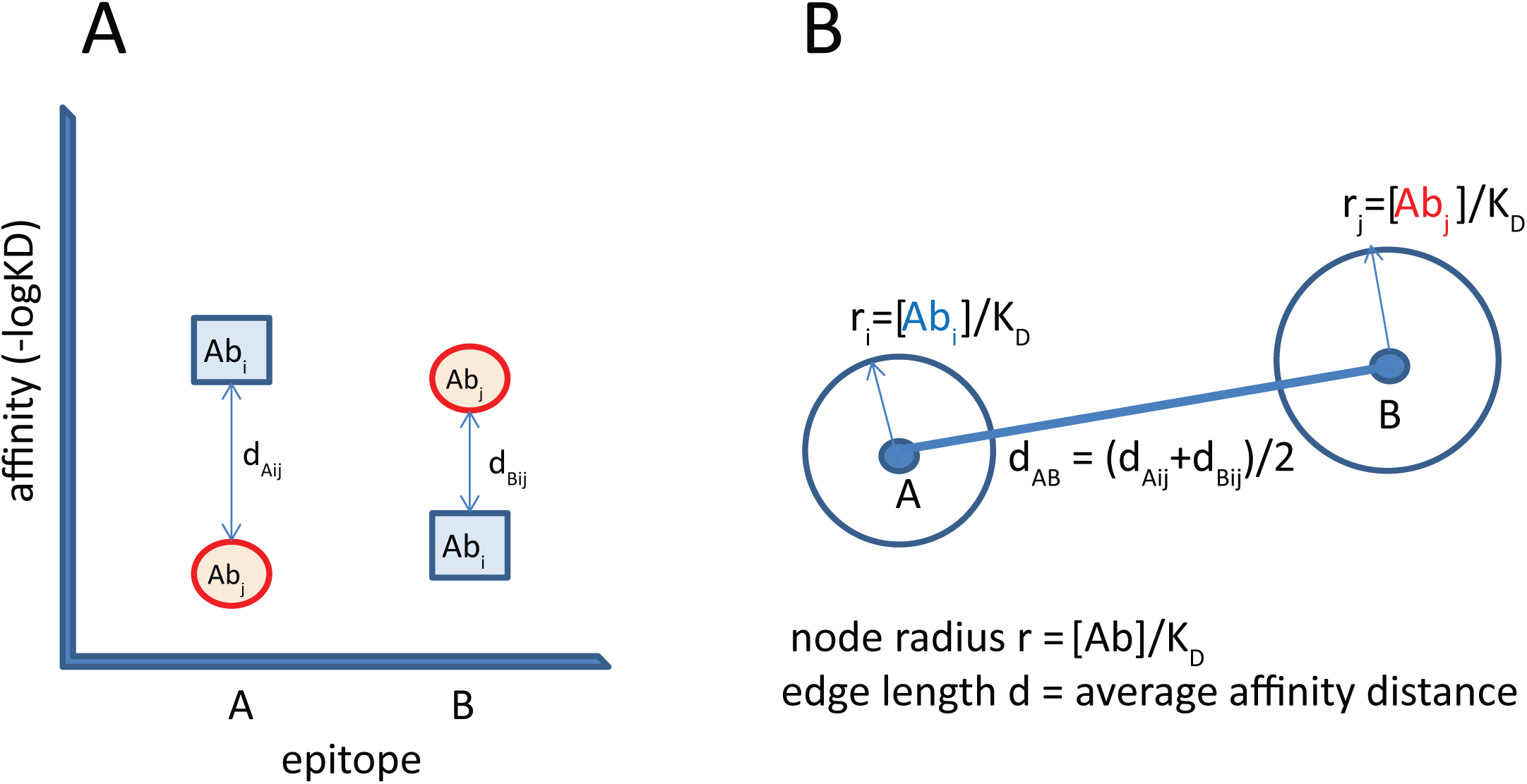
Definitions of the antibody network. Antibody Abi reacts with epitope A and cross-reacts with epitope B. Abj reacts with epitope A and cross-reacts with epitope B. Distances, expressed as negative logarithm of K_D_ are averaged to get the distance in the epitope space, providing an edge in the network. Free antibody concentration divided by K_D_ reflects antigen binding capacity. This value is displayed as the radius of the node. Changes in antigen concentration will trigger changes in node radius. New antigens entering the system may react with higher affinity with a given antibody and thereby change its position in the epitope space.

These points and spheres are connected by distances depicting cross-reactivity of the antibodies. We can characterize epitope similarity, or in other words antibody cross-reactivity, by averaging the differences in affinity between cognate antigen and the other antigen (Fig.5). As B cells are developing more and more points and spheres appear in the epitope space. Closely related antibodies, such the as progeny of a pre-BI cell, with identical heavy chains, will be close to each other and farther away from other clusters with a different heavy chain.

It is important to note that there is no absolute definition nor intrinsic property for a self or foreign epitope. Rather, the developing immune system carves out a space that will define the network of interactions defining self. We can call this space the extended immunological self (Fig.4). As IgM production increases these molecules can present as epitopes themselves, seeding new interactions and leading to the development of anti-idiotype antibodies^28^. The resulting closely knit network defines the space containing epitopes regarded as self. Once lymphoid organs are structured and the host is born, foreign molecules enter and various kinds of antibody responses develop. **TI responses increase Ab concentration without changing K_D_, thus the position of the node does not change only the size. TD responses also decrease K_D_, changing the position of a node in the epitope space and its distance to related clones**. Huge increases in affinity along with loss of polyspecificity result in the extrusion of the node from the network (Fig.6).

**Figure 6.**
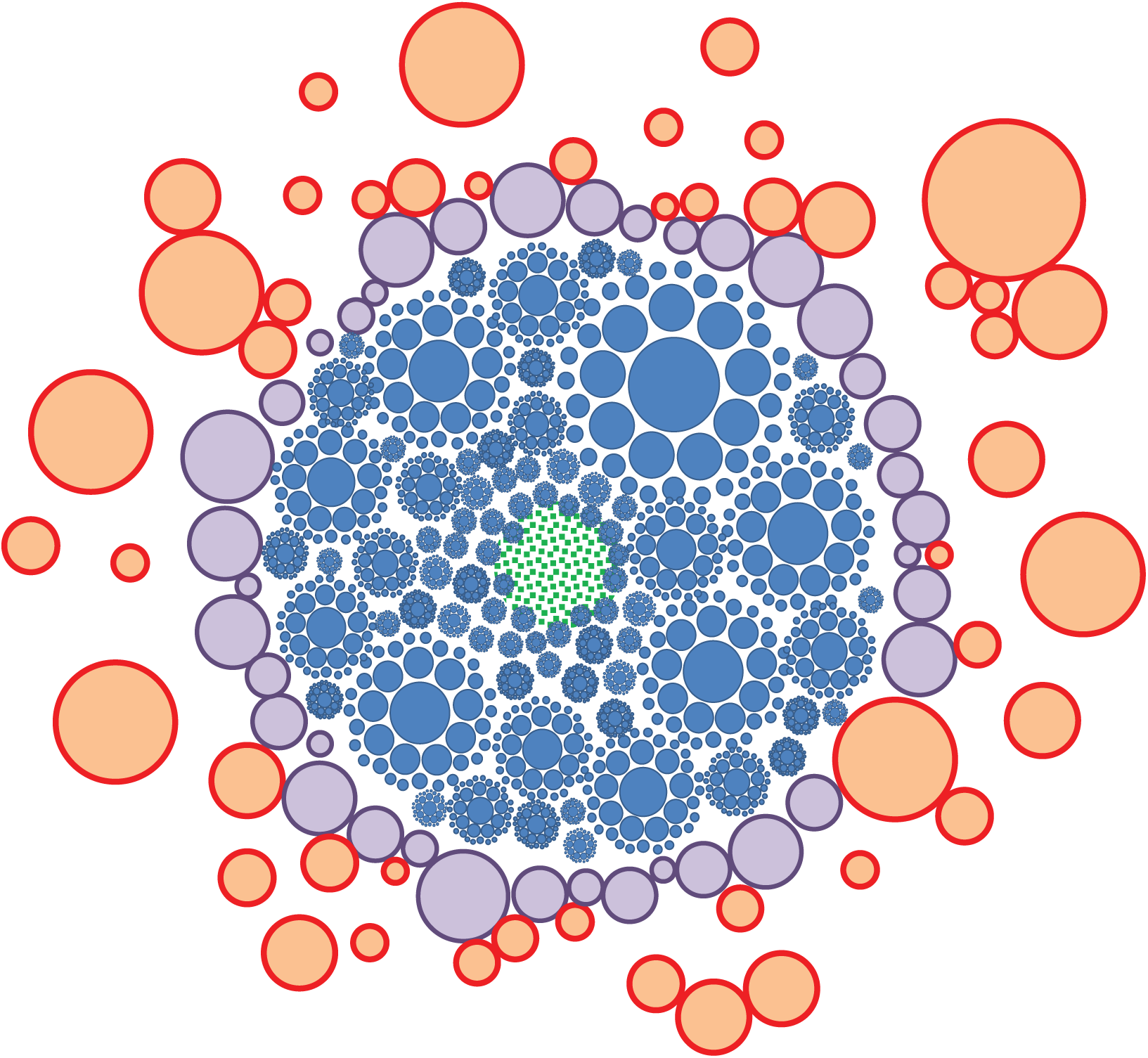
Simplified schematic representation of the antibody-epitome interaction network. Only nodes of the network are shown, edges are not drawn for the sake of simplicity but are represented as internodal distances. Blue nodes are natural antibody interactions, purple nodes are interactions with the accepted environment, red nodes are TD interactions in the epitope space. Green area represents B cells in the quiescent state, not producing antibodies but presenting themselves in the epitome space. Immunological self is a superconnected central component, while epitopes marked for elimination are extruded and disconnected.

The fact that an antigen reaches and stimulates a B-cell reflects that it has not been masked by antibodies from being recognized by that clone. Additionally, uptake via the BCR cn be accompanied by the uptake of lipids or proteins recognized by helper T-cells. These two events warn the immune system that a molecule that needs to be eliminated breached the network barrier. The production of high affinity antibodies means that a new node (connected to other clones with the same specificity but lower affinity) appears in the respective antigenic space, filing the space with an increased node size, which reflects increased [Ab]free/K_D_ value.

With the definitions we have built up

- B cells respond to [Ag] by developing into ASC
- B cells respond to [Ag] by changing KD
- [Ab] is regulated to reach K_D_ for control of [Ag]
- network node size is [Ab]/K_D_
- i-j internodal distance is |K_Di_-K_Dj_|
- in a network with N=number of B cells with distinct antigen binding properties nodes

we have outlines a dynamic, weighted network model of the humoral immune system, shown in a simplified two-dimensional representation in Figure 6. Because TI responses utilize polyclonal responses, antibody production against a given epitope region is always shared, [Ab]/K_D_ is always less than 1. For oligoclonal or monoclonal TD responses this value may exceed 1. The model shows the space filling property with multi-sized spheres, the core representing immunological self – a giant superconnected component of the network of antibodies.

## Concluding remarks

We have introduced homeostatic antibody model based on the assumption that the clonal humoral immune system seeks after an equilibrium between antibody and antigen. This requires that the membrane-bound form of antibody, the BCR, regulates the fate of cells that produce antibodies, as shown in our previous article. In this article we argued that both the affinity and the concentration of the antibodies produced in the host are tuned for silently degrading, carefully removing or aggressively eliminating antigen.

An important message of this approach is that molecular abundance is the defining factor for immunological self. Everything abundantly present in blood and in the bone marrow during the development of the system is self immunologically. Immunological homeostasis is about controlled catabolism and regeneration of self. Self is silently eliminated when ages and aggregates. Molecules binding with high affinity to cells screened for low-affinity self-binding and accompanied by danger signals are regarded as to be eliminated. In this sense low-affinity self-recognition is necessary not to avoid binding to self but rather to set a point of reference in the epitope space.

There have been several different approaches and theories to provide general, perhaps mathematical explanations for the complexity, the functioning of the immune system. The idiotype network theory^28^ originally worked out by Niels Jerne^29^, and the clonal selection theory elaborated by Frank MacFarlane Burnet^30^ identified different, seemingly contradictory concepts, which were resolved by Coutinho^31^ who suggested that “clonal as well as network organization co-exists in the immune system”. Our GQM is in agreement with this observation, as clonal responses generate a network, even though this network is a virtual antibody-antigen network of connectivity rather than a physical network. In Zvi Grossman’s horizontal networks^32^ “the relative probability of maturation increases with antigen dose and with affinity” and “proliferation can be uncoupled from differentiation under certain predictable conditions”, which are key assumptions in our model as well. Grignolio et al. recently compiled factors that potentially influence immunological identity, focusing on the changing nature of immunological self, which they conceptualize as “liquid self”^33^. The concept of immunological homunculus by Irun Cohen et al.^34^ states that non-reactivity can only create chaos, self-reactivity is required for control and order. Our model suggests that the need for self-reactivity is quite profane: controlled catabolism. As long as microbial epitopes are also efficiently cleared they may constitute part of the immunological self. The reactivity networks generated by Bransburg-Zabary et al.^35^ by antigen microarray analysis are probably the experimental observations closest to our model. In these experiments correlation analysis of antibody reactivity between individuals revealed the presence of connected specificities. The deconvolution of polyclonal antibody reactivity within individuals will be the key to generate experimental data directly supporting our model.

Quantitative, descriptive approaches to antibody networks currently rely on genetic information obtained by new generation sequencing technologies^36–38^. These can now generate antibody sequence repertoires with immense depth and fine resolution. While it is possible to sequence the B cell repertoire of a whole zebra fish^39^ we cannot do the same with a human B-cell repertoire. There will be a sampling bias depending on where the B cells are obtained from: bone marrow, blood, tonsils, lymph nodes, spleen. Additionally, sequencing cannot provide antibody concentrations, even less so free antibody concentrations or antigen concentrations in the sampled organism. Immunomics approaches will need to utilize databases on antibody-antigen interactions that have already been compiled (The immune epitope database, allergome database). The development of novel immunological methods is required for further formal proof of this theory and for its application in various fields of immunology. By combining genetic, immunologic and immunoinformatics data we expect that a true bioinformatics modeling of the complete human antibody network, its dynamics, disease-associated changes will be possible in the near future.

## Acknowledgments

This paper is dedicated to all immunology theorists, past and present, who advanced the field of immune network theories. I wish to thank Károly Liliom for consulting us on enzyme kinetics and Anita Orosz and Krisztián Papp for working with me on methodological approaches to quantitative immunomics. I thank my family and my colleagues for their patience and understanding in the moments when I was too immersed in thinking.

